# Ultrastructural and transcriptional changes during a giant virus infection of a green alga

**DOI:** 10.1101/2024.08.30.610430

**Authors:** Andrian P. Gajigan, Christopher R. Schvarcz, Cecilia Conaco, Kyle F. Edwards, Grieg F. Steward

**Affiliations:** Department of Oceanography, University of Hawai‘i at Mānoa, Honolulu, Hawai‘i 96822; Center for Microbial Oceanography Research and Education, University of Hawai‘i at Mānoa, Honolulu, Hawai‘i 96822; Marine Science Institute, University of the Philippines Diliman, Quezon City 1101, Philippines

**Keywords:** transcriptome, mimivirus, green algae, ultrastructure, *Tetraselmis*

## Abstract

The complete genome sequence of the *Oceanusvirus kaneohense* strain (Tetraselmis virus 1; TetV-1) was previously reported, but little is known about the virus infection cycle. Using a permissive *Tetraselmis* isolate (UHM1315), we estimated the eclipse period (4–8 hours), latent period (16 hrs), and burst size (800–1000) of the virus and documented ultrastructural and transcriptional changes in the host during infection. Putative viral factories and electron-dense inclusion bodies appeared in the cytoplasm of infected cells by 8 and 16 h post-infection, respectively. The nucleus and chloroplasts appeared to remain intact but reduced in size after 8 h. Transcriptome sequencing suggests that the viral genome codes for 830 transcripts. Those expressed early in infection (eclipse period at 0.25 and 4 hr) were related to the initiation of transcription, DNA synthesis, translation, and host immune repression. During the later, post-eclipse period (8, 12, 16 hr), virus structural genes were expressed. For the algal host, transcripts related to lipid metabolism and endocytosis were upregulated during the early phase, while those for protein modification/ turnover/ transport were downregulated. In the later period, host transcripts associated with basic cellular processes were upregulated, while genes related to morphogenesis/development were downregulated. Many of the most highly expressed virus and host genes were of unknown function, highlighting a need for additional functional studies.

## Introduction

Viruses in the phylum *Nucleocytoviricota*, the new taxonomic designation for the Nucleocytoplasmic Large DNA Viruses (NCLDVs), are characterized by their large double-stranded DNA genomes (up to 2.5 Mbp) and large virion sizes (up to 1500 nm) ^1–3^. The largest members in this phylum, often referred to colloquially as “giant viruses” ^4^, contain hundreds of genes that afford them a higher degree of autonomy compared to ‘smaller’ viruses ^5^, the smallest of which encode as few as two genes. In addition to a typical suite of DNA replication and transcription machinery, giant virus genomes may encode a diverse suite of products involved in metabolic functions previously only associated with cells. For example, some encode dozens of tRNAs and hundreds of components of the translation machinery ^6,7^. Others may encode rhodopsins ^8,9^, enzymes involved in the TCA cycle and glycolysis ^10,11^, histones ^12,13^, cytoskeletal proteins (actin, kinesin, myosin) ^14,15^ or, as was found for the virus studied here, TetV-1, enzymes involved in fermentation ^16^. In addition to the genes with diverse predicted functions, giant viruses also encode many ORFans ^17^ or genes with predicted amino acid sequences that are so divergent from any characterized protein that their functions remain a mystery. Even for the genes for which confident functions are assigned, how and when those genes are used by the virus is not always clear.

Transcriptional analysis complements genome sequencing by providing detailed insights into the operational program of a virus ^18^. Transcriptome sequences can reveal errors in genome sequencing and assembly ^17^ and improve genome annotations. Genes not predicted *ab initio*, for example, may be revealed by their presence in the transcript pool ^19–21^. Conversely, initially predicted genes may be absent from the transcriptome ^22^. Transcriptional analysis can also shed light on how gene expression is regulated, revealing, for example, changes in the promoter motifs used at different stages of infection ^19,23^, alternative splicing ^24,25^, transcription start and termination sites ^19^, and transcription termination mechanisms ^18,26^. Transcriptomics also allows for inferences about the interplay between virus and host metabolism ^23,27–29^. In some cases, host genes are upregulated that carry out functions not encoded by the virus ^30,31^. In other cases, the virus and host share homologous genes, but the version expressed (host vs virus) changes during infection ^27,32^.

Inferences about the timing of replication based on transcriptional patterns can be corroborated and augmented by observations of ultrastructural changes in infected cells. Electron microscopy of thin sections provides information on the mechanisms of virion entry ^33,34^, the location of replication and the formation of viral factories ^35^, the timing of the eclipse period, and other virus-induced changes to cellular ultrastructure ^7,32^. In this study, we investigated the infection cycle of the green alga-infecting Tetraselmis virus 1 (Tet-V1) using both electron microscopy and transcriptomics to (i) identify ultrastructural changes accompanying the infection progression, (ii) temporal expression of viral genes, and (iii) host responses to infection.

## Results

### TetV-1 life history traits and ultrastructure

In an initial experiment tracking the infection cycle of TetV-1, free virions remained consistently low for the first 15.5 hours post-infection (hpi), began a rapid increase by 16 hpi, and plateaued at 17–18 hpi (**Fig. 1**). Based on the number of new viruses produced and the host counts at the beginning of the infection, and assuming all cells are lysed, the burst size was estimated to be around 800–1000.

**Figure 1.**
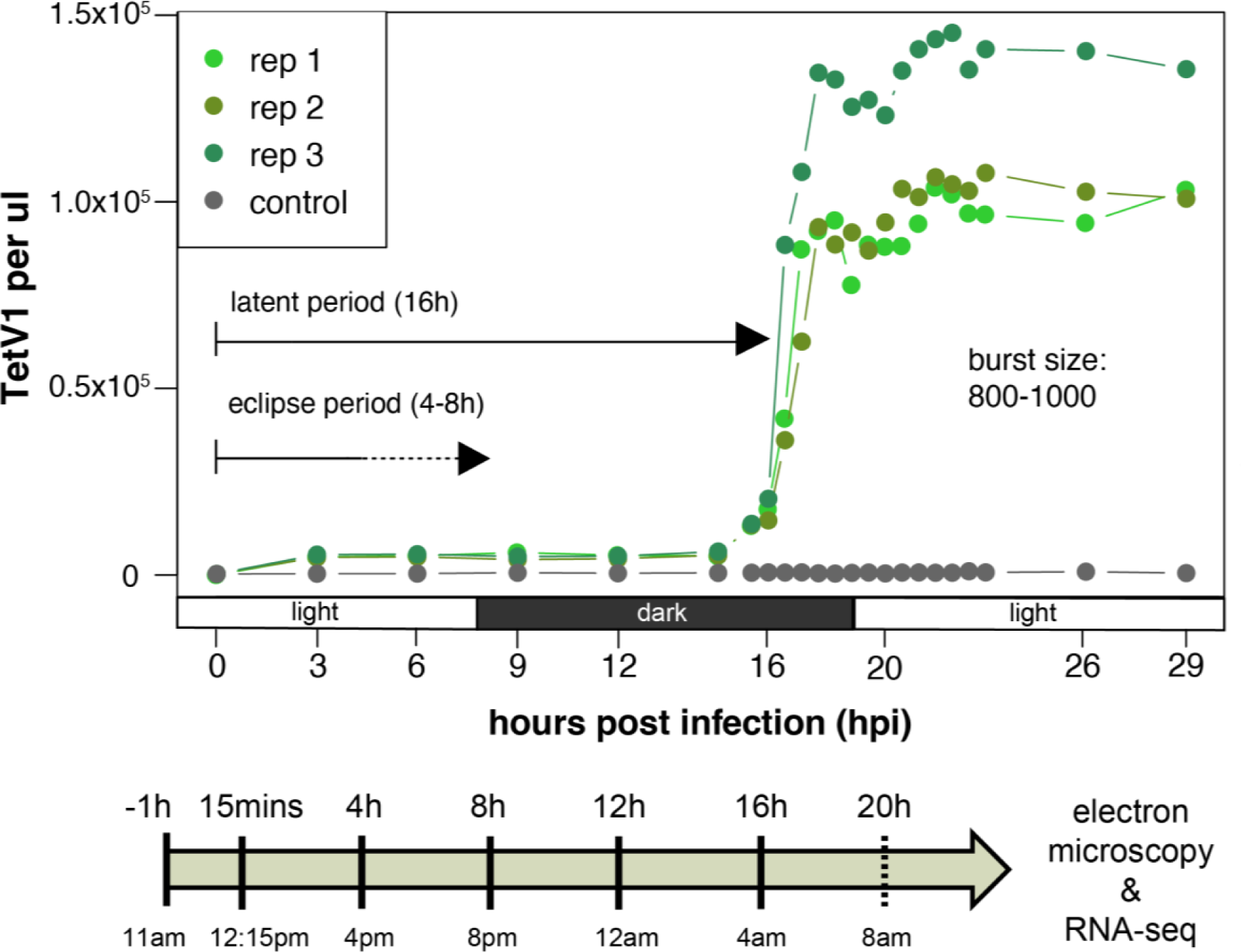
Tetraselmis-TetV1 infection cycle. Virus counts were measured using flow cytometry. Subsequently, estimates of the latent period and burst size were determined. Eclipse period was determined using electron micrographs. Samples for electron microscopy and RNA sequencing were obtained throughout the time series except at 20 hpi (solid lines on the arrow below the graph), where RNA sequencing was not implemented.

Examination of thin sections of uninfected algal cells by electron microscopy revealed the intact organelles expected in a healthy cell such as the nucleus, nucleolus, chloroplasts, mitochondria, vacuoles, pyrenoids, and an eyespot (**Fig. 2A**). Four basal bodies (base of flagella) are also evident (**Supp Fig. 1**). For infected cells, the initial attachment and entry process of the virus was not observed in any of the examined cells, likely because of the low temporal resolution of sampling. Cells at 15 minutes and 4 hpi were similar to the uninfected cells, with no apparent changes to subcellular structures or evidence of virion assembly (**Fig. 2B-C**). Intracellular virions were first observed in samples taken at 8 hpi, which puts the eclipse period somewhere between 4 and 8 hpi (**Fig. 2D**). By 8 hpi, there was also a change in the fine-scale structure of the cytoplasm. At this point, organelles appeared to be compressed around the periphery of the cell, the expanded area of cytoplasm was more granular, and putative viral factories (VFs) were present. The VFs are localized areas within the cytoplasm with incomplete capsids near the center and complete capsids further out. VFs were present for the remainder of the latent period, (**Fig. 2E-F),** but multiple, well-defined, darker staining inclusion bodies (IBs) also appeared in the cytoplasm by 16 hpi (**Fig. 2F**). At this point most cells were filled with virions, anticipating the imminent lysis, which is evident from the rapid increase in free virions over the next two hours (**Fig. 1**). Some intact cells at the late stages of infection with both VF and IB were still observed at 20 hpi (**Fig. 2G-I**), either because of a non-synchronous start of infection, or variation in latent period length. Higher magnification views of one such cell illustrate the incomplete capsid structure at the core of the VF and a darker staining edge of the IB, suggesting a membrane-bound compartment. (**Fig. 2G-I**).

**Figure 2.**
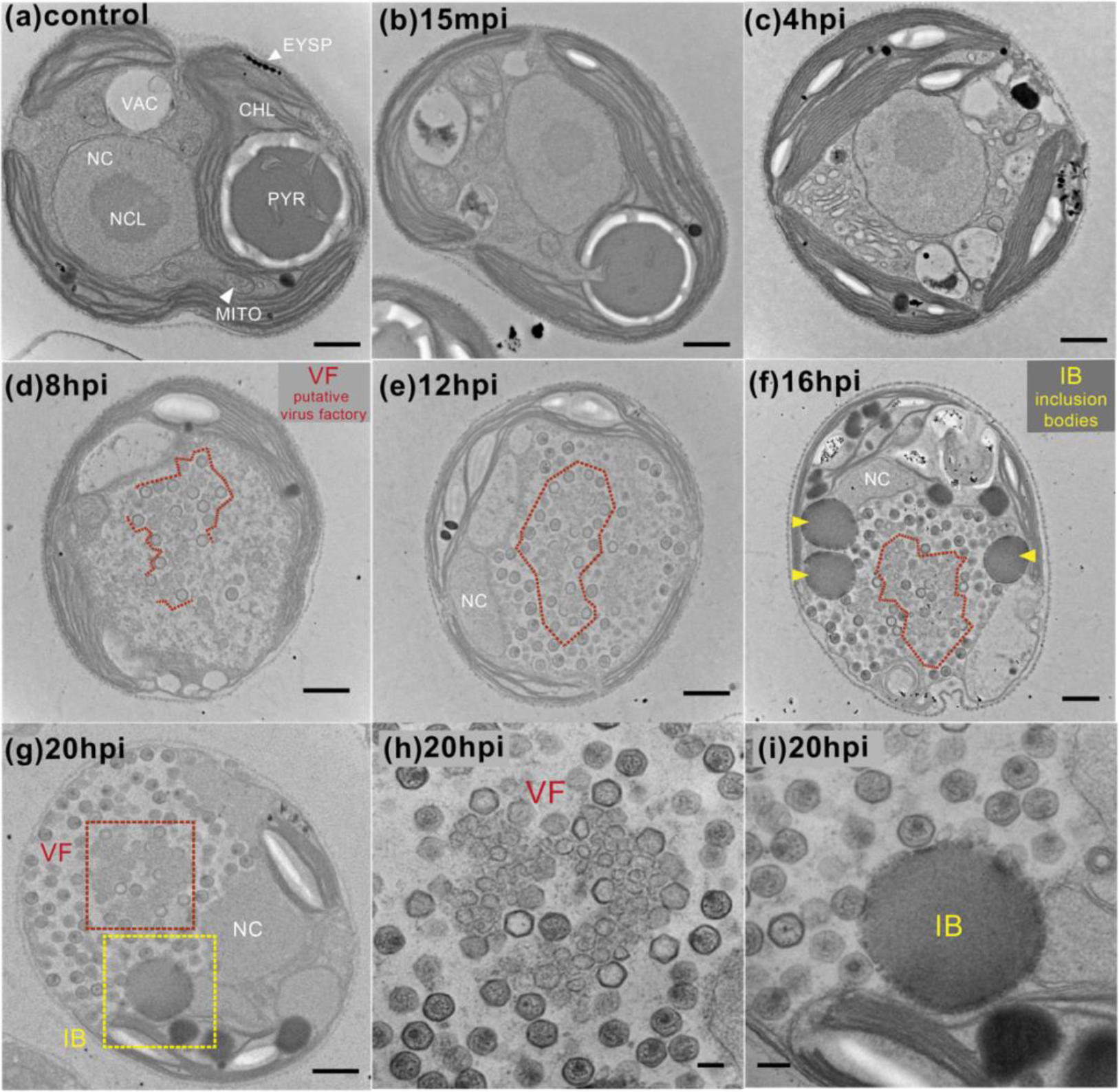
Ultrastructural changes of *Tetraselmis*-TetV1 virocells across infection time course. (**a**) Apparent in the uninfected algae are typical organelles such as the nucleus (NC), nucleolus (NCL), chloroplast (CHL), vacuoles (VAC), mitochondrion (MITO), as well as algae-specific photoreceptive organelles (eyespot; EYSP) and CO_2_-fixation center (pyrenoid; PYR). (**b, c**) No changes were observed in the early stages of infection (15 mins and 4 hpi). (**c, d**) Further inspection of electron micrographs showed the first appearance of intracellular virions at 8 hpi but not 4 hpi, indicating an eclipse period between 4 and 8 hpi. (**e, f, g**) Representative images of the virocell close to the latent period show the presence of (**g, h**) a putative virus factory (VF) and (**i**) inclusion bodies (IB). Scale bar = 800 µm (a–g) or 200 µm (h, i).

### TetV-1 transcriptome

The TetV-1 genome was sequenced previously and shown to contain 663 genes predicted using Prodigal ^16^. The current RNA-seq pipeline detected 830 transcripts, of which 167 (20%) are multi-exon. Included in these are 292 transcripts that are longer than the Prodigal-predicted genes, 27 that contain overlapping exons with Prodigal-predicted genes, one transcript that is within an intron, and 10 transcripts contain exons that overlap with Prodigal-predicted genes but are on the opposite strand.

The relative abundance of reads that map to TetV-1 steadily increased early in the infection and peaked at 8 hpi, at which point they accounted for ∼58% of the total reads (**Supp Fig. 2A**). The proportion of viral reads subsequently decreased, presumably when transcription ceases and other processes, such as DNA synthesis, packaging, and assembly predominate until lysis at around 16 hpi (**Supp Fig. 2A**). The average RNA yield remained constant in the control sample, while it seems to decrease in the infected samples (**Supp Fig. 2B**). Global clustering of TetV-1 gene expression profiles resulted in the earliest time points (15 mpi and 4 hpi) forming one cluster and the three later time points (8–16 hpi) forming another (**Supp Fig. 3**). Given this pattern and the limited temporal resolution of the sampling, we labeled genes in subsequent analyses as either “early” or “late,” depending on when they were most highly expressed. These categories are equivalent to eclipse and post-eclipse periods. After a filtering step to remove transcripts that were relatively constant (those with a variance less than one across samples), 94% of the viral genes (782 out of 830) were retained. Nine of the top ten highly expressed genes are of unknown function (TetV_630, 071, 278, 511, 015, 023, 024, 013, and 405) (**Fig. 3**).

**Figure 3.**
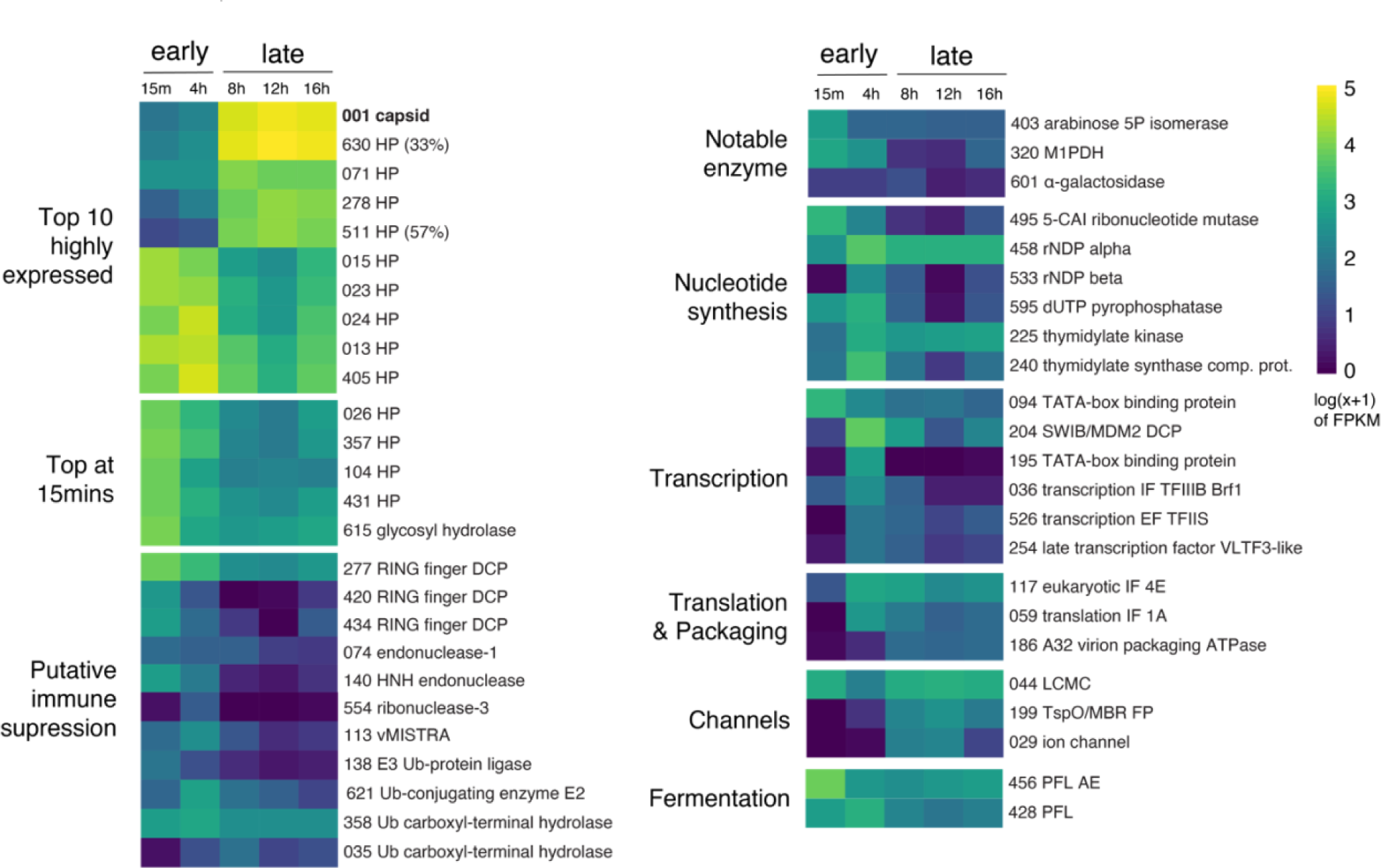
Temporal expression of select TetV1 genes. Genes are grouped based on their function, as described above. HP = hypothetical protein (percentage shows the relative abundance in proteome with respect to capsid), DCP = domain-containing protein, vMiSTRA = TetV-1 viral mitochondria stress response algae, Ub = Ubiquitin, M1DPH = mannitol 1-phosphate dehydrogenase, 5-CAI = 5-(carboxyamino)imidazole, rNDP = ribonucleoside-diphosphate reductase, IF = initiation factor, EF = elongation factor, LCMC = large conductance mechanosensitive channel, FP = family protein, PFL = pyruvate formate lyase, PFL AE = pyruvate formate lyase 1 activating enzyme, FPKM = Fragments Per Kilobase of transcript per Million mapped reads.

### ***A.*** Early infection stage

Expression of viral transcripts was detected at least as early as 15 min after infection (**Fig. 3, Supp Fig. 4, Supp Table 1**). Four highly expressed transcripts at the 15 mins post infection have no known or predicted function (TetV_026, 357, 104, and 431).

**Figure 4.**
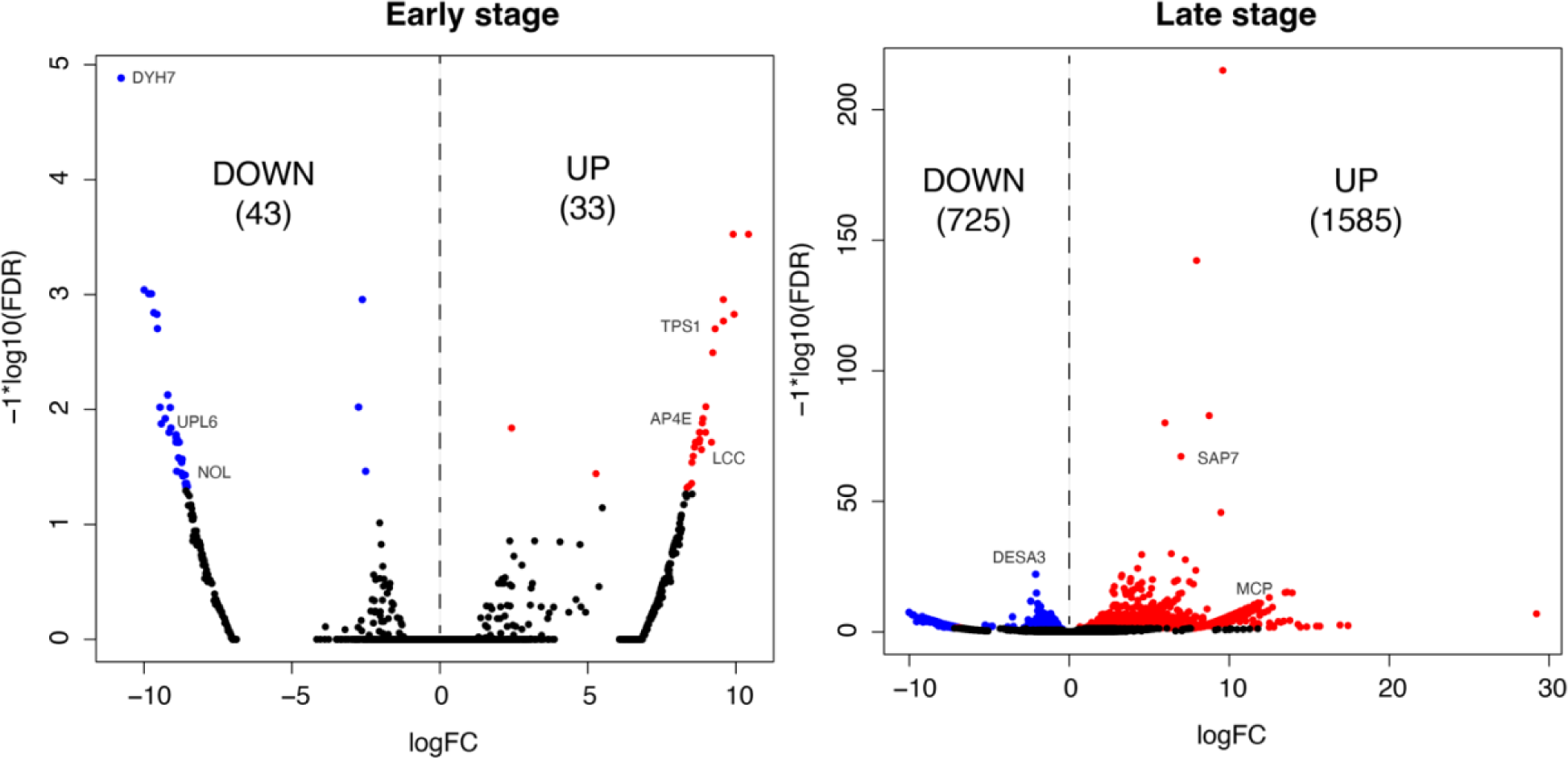
Differentially expressed (DE) host genes during the early and late stages of the infection. The number of down or upregulated genes are shown in parentheses. Colored points represent significant genes at FDR (false discovery rate) ≤ 0.05. FC=fold change. 0.25 and 4 hpi represent early-stage samples while 8 and 12 hpi represent late-stage samples.

Several enzymes involved in carbohydrate metabolism are expressed early such as alpha-galactosidase (TetV_601), glycosyl hydrolase (TetV_615), arabinose 5-phosphate isomerase (TetV_403) and mannitol 1-phosphate dehydrogenase (M1DPH, TetV_320) (**Fig. 3**). A putative host immune suppression gene, vMISTRA (TetV_113), and various RING domain-containing proteins (TetV_277, 420, 434) that are implicated in apoptosis inhibition are also expressed early as is the virus-encoded ribonuclease-3 gene (TetV_554) (**Fig. 3**). Most ubiquitination genes are expressed throughout the infection (TetV_035, 358, 621) except one that seems to be expressed early, E3 ubiquitin-protein ligase MIEL-1 like protein (TetV_138). The virus-encoded fermentation genes, pyruvate formate-lyase (vPFL; TetV_428) and pyruvate formate lyase activating enzyme (vPFLA; TetV_456) are somewhat higher in the early phase than in late stages, but were essentially constitutively expressed (**Fig. 3**).

Expression of genes involved in transcription began in the early stage and expression of some was sustained throughout the infection (**Fig. 3, Supp Fig. 4**). Transcription factors such as transcription initiation factor TFIIIB Brf1 (TetV_036), transcription activator SWIB/MDM2 (TetV_204), and TATA-box binding proteins (TetV_094 and 195) were among those expressed early. In addition, transcription elongation factor TFIIS (TetV_526) and late transcription factor VLTF3-like (TetV_254) were expressed early and were sustained throughout the infection (**Fig. 3**). Similarly, DNA-directed RNA polymerase subunits were expressed early and sustained in the late stage. These included Rpb2 (TetV_196, 616), Rpb3/Rpb11 (TetV_250), Rpb5 (TetV_078), Rpb6 (TetV_177), Rpb7 (TetV_286), and Rpb10/RpoN (TetV_156) (**Supp Fig. 4**). Expression of mRNA capping enzymes (TetV_090 and 219) also began in the early stage (**Supp Fig. 4**).

Early genes involved in nucleotide synthesis included ribonucleoside-diphosphate reductase subunit alpha (TetV_458) and beta (TetV_533), thymidylate kinase (TetV_225), thymidylate synthase complementing protein (TetV_240), 5-(carboxyamino) imidazole ribonucleotide mutase (TetV_495), and dUTP pyrophosphatase (TetV_595) (**Fig. 3**). Early genes involved in DNA replication, synthesis and repair genes included DNA polymerase B (TetV_567), DNA topoisomerase 2 (TetV_404), and DNA clamp, proliferating cell nuclear antigen (TetV_607) (**Supp Fig. 4**). Genes involved in DNA repair (DNA repair exonuclease SbcCD, TetV_472; DNA repair ATPase SbcC, TetV_662) appeared early, as did others that remained relatively high through the late stages such as DNA mismatch repair ATPase MutS (TetV_243) and deoxyribodipyrimidine photolyases (TetV_298 and 522) (**Supp Fig. 4**). Finally, initiation of translation also starts at the early stages of infection based on the expression of eukaryotic initiation factor 4E (TetV_117) and translation initiation factor 1A (TetV_059) (**Fig. 2**).

### ***B.*** Late infection stage

Genes encoding proteins related to virion structure and packaging, such as the major capsid protein (TetV_001) and VV A32-like virion packaging ATPase (TetV_186) (**Fig. 3**) were highly expressed in late stage. Genes encoding a functional potassium ion channel (TetV_029) ^36^, a large-conductance mechanosensitive channel (TetV_044), and a transmembrane protein/ion channel (TetV_199) were also expressed in late stages (**Fig. 3**). Genes encoding some enzymes were also expressed in the late stage such as glycosyltransferases (TetV_241 and 318), sulfatases (TetV_008 and 050), lipid catabolic enzymes, lipase (TetV_056) and phosphatase (TetV_413), proteases (TetV_461 and 556), and redox enzymes (TetV_019, 191, 236, and 626) (**Supp Fig. 4**). Different methylation genes appear to be expressed at distinct stages of the infection. There is one methylation gene expressed early, the SET domain-containing protein (TetV_181), whereas four methyltransferases are expressed in the late stage (TetV_211, 484, 501, 608) (**Supp Fig. 4**).

### Tetraselmis sp. (host) response

Before the host transcriptome assembly, TetV-1 reads were removed, leaving 308 million paired-end reads. Trinity *de novo* assembly yielded 159,006 transcripts corresponding to 69,817 genes (**Supp Table 2**). The assembly is of high quality based on metrics such as N50 (3,244 bp), read alignment (99%), and the presence of 96% of Chlorophyta single copy genes (**Supp Table 2**). Two additional filtering steps were implemented: First, only transcripts with enough read support according to TransRate were retained, leaving 91% of the genes. TransRate calculates contig scores based on read mapping evidence, identifying contigs that are chimeric or misassembles, among others. Second, only the highest expressed isoform per gene was retained for downstream analysis (**Supp Table 3**). Of the remaining 63,860 genes, 83.7% are expressed at > 1 TPM (transcripts per million transcripts), while only 37% are expressed at > 5 TPM. Around 14-15% of the genes have BLAST/HMMER matches to SwissProt Uniprot and Pfam databases (**Supp Fig. 5**). Out of the annotated genes, 46.4% are closely matching eukaryotic genes (the majority being Viridiplantae homologs), while the rest are closely matching Bacteria (51.3%), Archaea (1.6%), and viral (0.6%) genes (**Supp Fig. 5**).

**Figure 5.**
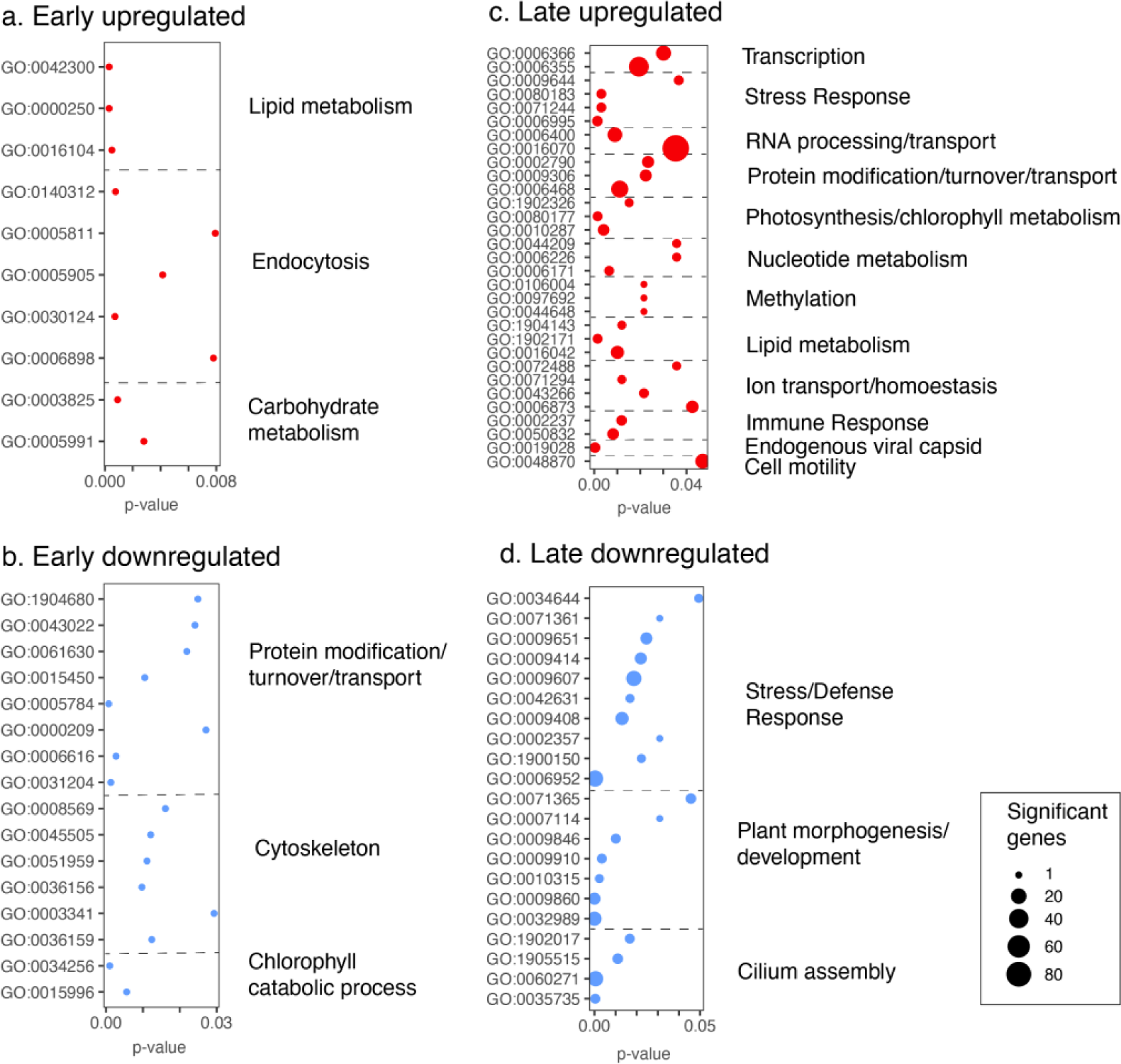
Tetraselmis sp. (host) response. Enriched GO (gene ontology) terms illustrating the functions that were either upregulated or downregulated during the early and late stages of the infection

We found seven host transcripts related to fermentation with homology to pyruvate formate lyase activating enzyme (PFLA), but no transcripts related to pyruvate formate lyase (PFL) (**Supp Fig. 6**). Four host transcripts matched to formate acetyltransferase (PFLB) (not shown). Only one of the seven host PFLA genes showed significant expression, and, like the viral-encoded PFLA, the host PFLA seems to be expressed in high levels early in the infection.

To determine the host response to infection, differentially expressed genes were determined at early (15 mpi, 4 hpi) and late (8 hpi, 12 hpi) stages of infection (**Fig. 4**). The control sample corresponding to the 16 hpi sample was potentially cross-contaminated with viral reads so, this time point was excluded from analysis (**Supp Fig. 2**). A gene ontology enrichment test was performed to identify overrepresented functions in the differentially expressed genes. Few differentially expressed genes were observed during the early stage (33 upregulated and 43 downregulated out of 63,860 genes). The upregulated gene set shows enriched GO terms for carbohydrate and lipid metabolism and endocytosis (mainly attributable to adaptor protein complex 4; AP4E gene) (**Fig. 4, Supp Fig 5A, Supp Table 4**). The downregulated gene set includes GO terms related to protein modification/transport/turnover (ex. UPL6, E3 ubiquitin-protein ligase; SC61A, transport protein Sec61 subunit), cytoskeleton (ex. Dynein heavy chain 7; DYH7) and chlorophyll catabolic process (ex. NOL, Chlorophyll(ide) b reductase) (**Fig. 4, Supp Fig 5B, Supp Table 5**).

A much greater proportion of genes were differentially expressed in the late stage (1,585 upregulated and 725 downregulated) (**Fig. 4**). Among the genes upregulated were those with GO terms indicating involvement in immune and stress response, nucleotide, and lipid metabolism, methylation, RNA processing and transcription, ion transport, protein modification, and cell motility (**Fig. 5C, Supp Table 6**). Interestingly, putative endogenous major capsid proteins were also upregulated in the *Tetraselmis* host (**Fig. 4-5, Supp Fig 7**). In the downregulated gene set, enriched GO terms included stress/defense response, “plant-related morphogenesis,” and “cilium assembly” (**Fig. 5D, Supp Table 7**).

Differentially expressed genes and enriched GO terms in samples representing light (11 am, 12:15 pm, 4 pm) vs. dark (8 pm, 12 am) cycles were determined in the uninfected (control) time series (**Supp Fig. 8**). As expected, enriched GO terms in the *Tetraselmis* light cycle include photosynthesis, thylakoid, and chloroplast, among others, while in the dark cycle, DNA and nuclear processes are enriched (**Supp Fig. 9**).

## Discussion

### TetV-1 life history and ultrastructural changes

With this study, we provide details of the life cycle of TetV-1 when infecting a marine coastal strain of the green alga *Tetraselmis* sp. (UHM1315). Latent period and burst size are among the most important to parameterize when modeling the spread of viral infections through phytoplankton populations ^37^. However, they are not fixed traits. A virus that infects a cell under sub-optimal conditions (e.g., growth-limiting temperature, irradiance, or nutrient supply) may present a significantly longer latent period and reduced burst size ^38^. Despite the context-dependence of these traits, our estimates for TetV-1 were very similar to those reported for its closest cultivated relative Pyramimonas orientalis virus, or PoV-01B. Like TetV-1, PoV-01B is a green-alga infecting virus in the family *Allomimiviridae*. The latent period (14–19 h) and burst size (800-1000) reported for PoV-01B ^39^ were indistinguishable from our findings for TetV-1, despite the former being assayed at lower temperature (15 °C) and irradiance (40 µmol photons per m^-2^ s^-1^). Undoubtedly these values would change for either virus under more stringent growth limitations, but our objective in determining latent period in this study was primarily to determine the endpoint of the virus life cycle as context for interpreting changes in ultrastructure and gene expression under the same conditions. Under our controlled conditions, the latent period accounted for more than a quarter, but less than half of the latent period and provided a convenient visual break point for early (replication) and late (assembly) stages of infection.

The appearance of viral factories in the cytoplasm and the persistent integrity of the nucleus during infection is similar to observations for some members of the families *Mimiviridae* and *Poxviridae* ^35,40^. The reduction in *Tetraselmis* chloroplast size during TetV-1 infection is similar to observations in *Emiliania huxleyi* when infected with EhV, but in the latter case, the nucleus was degraded ^32^. The putative VFs produced by TetV-1 are similar to structures that also have been referred to variously as viroplasm, replication organelles, assembly sites, or inclusion bodies in other virus-host systems ^41–44^. VFs concentrate viral replication machinery ^45^, which is proposed to protect viruses from host defenses ^41,44^. VFs are observed in other giant viruses, such as ApMV ^46^ and Tupanvirus ^47^. In ApMV, the VF is the source of viral DNA and is associated with empty virus capsids ^48^. The vesicular or angular shapes of apparent capsid components seen in TetV-1 virus factories are also present in PBCV-1 ^49^. In addition to the VFs, we observed other structures we refer to generically as inclusion bodies, which is a broad term for structures that appear in most virus-infected cells, but little is known about them ^44^. It remains to be seen if the inclusion bodies play a specific role in TetV-1 replication, or they are simply a cytopathic byproduct of the significant metabolic disruption caused by infections.

### TetV-1 transcriptional landscape

Transcription is the first major biosynthetic step after infection with a dsDNA virus. Immediate transcription is not unusual for giant viruses, with transcription happening within minutes in some cases ^28,50^. Transcription can even initiate within the virion prior to DNA unpackaging in the cytoplasm ^18^. Rapid transcription is possible because transcriptional proteins are packaged within the virion^18,51,52^. Like other NCLDVs such as AaV, ApMV, PBCV-1, and CroV, all known TetV-1 genes, as well as the 167 multi-exon transcripts discovered here, are expressed at least once in the infection cycle. In TetV-1, tRNAs were also detected by a sequencing library preparation method that enriched poly-A transcripts. This suggests that, unlike eukaryotic cellular tRNAs, those in TetV-1 are polyadenylated, as has been reported for tRNAs of other mimivirus ^53^. This suggests that RNA polymerase II is used for transcribing both tRNAs and protein-coding genes.

It has been hypothesized that the presence of exons and introns in viruses can disrupt nuclease cleavage sites ^54^ thereby providing protection from host nucleases, and this may be the case for giant viruses. Another possibility is that splicing does not benefit the virus but is a remnant of horizontally transferred genes that host spliceosomes recognize. Regardless of the selective forces that lead to their presence, the existence of multi-exon transcripts and alternatively spliced genes in TetV-1, and previous studies highlighting the rich alternative splicing repertoire of NCLDVs ^20,24,25^, emphasize that gene annotations based solely on genome sequences miss much of the complexity that underlies viral replication.

As has been found in transcriptional analyses of other giant viruses, most of the genes showing the highest expression are of unknown function ^22,29,30,55^. For those with a predicted function, the functional categories expressed in the loosely defined ‘early’ and ‘late’ stages of the TetV-1 infection were similar to those reported at similar stages of infection for many different NCLDVs^18,20,23,26–30,55–58^.

### Early-stage viral genes

Viral ‘early’ genes are typically initiation factors for DNA replication/transcription/translation, immune evasion, or nucleotide metabolism genes, as was the case for TetV-1. Also apparent at the early stages was the expression of enzymes that are likely involved in the destruction/degradation of host cell wall (alpha-galactosidase, TetV_601; glycosyl hydrolase, TetV_615), virion-cell recognition (arabinose 5-phosphate isomerase, TetV_403) or controlling intracellular osmolarity (mannitol 1-phosphate dehydrogenase M1PHD, TetV_320) ^59^ (**Fig. 2**). Mannitol has also been shown to suppress ROS-based defense (reactive oxygen species) against pathogens ^60^, thus it is possible that the virus is increasing the mannitol pool to counter host defenses. The early expression of nucleotide metabolism genes such as ribonucleotide reductase, rNDP (TetV_458, 533) and thymidylate kinase (TetV_225) and synthase complementing protein (TetV_240), is likely a response to the increased nucleotide demands of the virocell.

Other ‘early’ genes may be involved in evading the host’s innate immune response. The putative immune suppression gene, vMISTRA (<u>v</u>iral homolog of <u>MI</u>tochondria <u>ST</u>ress <u>R</u>esponse <u>A</u>lgae, TetV_113), is hypothesized to counteract viral-induced apoptosis ^61^. In addition, RNAse III (ribonuclease III) is known for its role in gene silencing with the potential to shift virocell global gene expression. Ubiquitination proteins (TetV_035, 358, 621) and other nucleases may play a role in keeping host proteins and DNA at bay (**Fig. 2**). Various RING domain containing proteins implicated in apoptosis inhibition, are also expressed early (TetV_27, 420, 434) (**Fig. 2**). This is the case for the Ascovirus IAPs (<u>I</u>nhibitors of <u>Ap</u>optosis) gene which encodes an E3 ubiquitin ligase with RING domain ^58^. TetV-1 encodes multiple copies of RING domain-containing genes that might represent gene duplications. Expansions of viral genetic content through gene duplication (i.e., a “gene accordion”) were suggested to be a mechanism by which poxviruses adapt to host antiviral defenses as new, duplicated genes can freely acquire mutations that facilitate virus evasion of the cellular immune system ^62^.

The highly expressed genes in the early stage of infection are located near each other (TetV_013, 015, 023, and 025) (**Fig. 2**). Spatial clustering of genes that show similar temporal expression suggests that a nucleoprotein/histone-like structure or other epigenetic modification (ex. methylation or acetylation) might govern their coordinated expression. Shared regulatory sequences (e.g., a promoter or enhancer) or operon-like structuring that enables synchronized gene expression can also explain this spatial clustering of genes. Spatial viral genome regulation is observed in EhV wherein a specific subregion (EhV218-366) is highly expressed at 1 hpi ^32^, and in PBCV-1, the top 50 expressed genes at 7 mins post-infection are also co-located. However, spatial clustering and temporally coordinated expression is not observed in other NCLDVs, such as the Marseillevirus ^29^.

### Late-stage viral genes

While the early phase of infection can be characterized as the preparation stage for viral production, the late stage is characterized by transcripts suggesting sustained biosynthesis of materials needed for viral assembly and egress. The late-stage transcripts are structural proteins, DNA packaging materials, transcripts/proteins used for egress, or transcripts/proteins that may be packaged in virus particles, among others.

Strikingly, most of the highly expressed late genes have unknown functions. These hypothetical genes/proteins that are co-expressed with the major capsid protein in high relative abundance in both transcriptome and virion proteome (TetV_511 and 630) are likely structural and excellent candidates for further functional characterization (**Fig. 2, Supp Table 1**) ^63^. It is also apparent that there are genes with unknown functions that display low to moderate gene expression (**Supp Table 1**) but are found in TetV-1 virion proteome such as TetV_155, 247, 258, 362, 378, 412, 463, 481, and 597 ^63^. This could also be subject to further functional screening. Various transmembrane proteins and ion channels are also expressed late (TetV_029, 044, 199) (**Fig. 2**). These proteins are likely involved in controlling cellular ion concentration or pH and could be involved in either virus entry or exit, depending on whether they are used in late-stage infection or packaged inside the virions. If a given transmembrane protein is used in the late stage, its function might be to alter cell osmotic pressure, facilitating exit. If packaged inside the virions, an ion channel may act at the early stage of infection in a manner similar to that observed for PBCV-1, where the potassium ion channel triggers depolarization and loss of ions, resulting in lower cellular pressure and triggering injection of viral DNA into the host ^64,65^. Here, among the three transmembrane proteins/ion channels, only the large conductance mechanosensitive channel (TetV_044) seems to be packaged inside the virions, based on virion proteome ^63^. It could also be possible that ion channels are crucial in controlling a balance of intracellular ions favorable for viral protein activity or mediating calcium/potassium-dependent signaling ^36,66^.

The expression of thioredoxin in the late stage (**Supp Fig. 4**) can be attributed to its role in regulating the cell redox state associated with cell death ^67,68^. In addition, late-stage methyltransferases (TetV_211, 484, 501, 608) might be responsible for the methylation of newly synthesized viral DNA (**Supp Fig. 4**). The sulfatase (TetV_008) that is highly expressed as transcripts in the late stage and has a high proteome signal in the virion can suggest the packaging of this specific enzyme (**Supp Fig. 4**). This enzyme can play a role in host surface or cytoplasm remodeling, as it catabolizes sulfated algal polysaccharides and glycolipids ^69^.

Here, we used conventional bulk RNA-seq which can be considered as the average state of virocell transcriptome. Because not all cells are infected at precisely the same time, there is some loss of resolution in the timing of expression when bulk sampling, as illustrated by single-cell transcriptomics of *E. huxleyi*-EhV ^23^. Furthermore, other RNA sequencing approaches could offer additional insights. For instance, CAGE-seq and 3’ RNA-seq will be required to ascertain transcriptional start and termination sites ^19^. Despite these limitations, our transcriptome analysis allowed the delineation of gene boundaries, identification of spliced transcripts, and determination of TetV-1 genes and their temporal profiles.

### Host metabolic reprogramming

This study demonstrated host responses to infection by analyzing differential gene expression and gene ontology enrichment analysis. These analyses paint a broad picture of host metabolic reprogramming in response to the virus attack. Host reprogramming can be accomplished through various mechanisms, such as viral-encoded nucleases that preferentially degrade host mRNAs, viral methylases and epigenetic modification systems, and virus-encoded transcriptional and post-transcriptional regulatory controls that affect host gene expression, among others. Here, we cannot ascertain the mechanism of host reprogramming, but nonetheless observed relative up-or downregulation of host genes as a response to giant virus infection.

In this study, few genes showed significant changes in expression during the early stage (76 genes out of 63,860 *Tetraselmis* genes). However, a host gene related to endocytosis, adaptor protein complex 4 subunit epsilon (AP4E), was upregulated. Adaptor protein complexes are host factors that interact with viruses to facilitate entry [reviewed in ^70^]. A related NCLDV, Vaccinia virus, for example, was shown to interact with its host’s adaptor protein complex 2 (AP2) ^71^. This circumstantial evidence suggests that AP4 may act as a critical host factor for TetV1 infection, although confirmatory tests are needed. A relatively higher number of host genes (2,310) are differentially expressed in the late stage. During this stage, we observed extensive changes in the ultrastructure, so, unsurprisingly, many cellular processes are disrupted relative to the control. Putative endogenous MCPs are upregulated at the late stage, which agrees with a recent survey that shows that protists, including various *Tetraselmi*s species, express endogenous MCPs ^72^. We are unsure if these are functional capsids or are degraded or repurposed genes that retained their transcriptional signals and are thus expressed upon infection. The lipid metabolism genes upregulated in both early and late stages are presumably associated with cytoplasmic rearrangement, creating lipid-rich virus factories ^42^. Downregulated host genes across the infection include those associated with the cytoskeleton, cilium/flagellar assembly, morphogenesis, and development, which seems consistent with disruption of cellular structure in favor of creating virus factories. It is worth noting, the constitutive expression of both viral and host-encoded fermentation genes (PFL and PLFA). Host pyruvate formate-lyase (PFL) was not detected in the transcriptome; however, *Tetraselmis striata,* the only *Tetraselmis* species with a published genome, contains a PFL (Protein ID: 421160; TSEL_012203.t1) ^73^. Hence, it is possible that PFL is also present in *Tetraselmis* sp. KB-FL45 but is below the detection limit of our sequencing approach.

## Conclusion

By tracking changes in the transcriptional activity and ultrastructure of a green alga infected with an allomimivirus, several broad patterns emerged consistent with responses to infection seen in other NCLDVs and viruses in general. Specifically, viral genes expressed early in infection were those involved with carbohydrate degradation, immune evasion, transcription initiation, and nucleotide metabolism, among others. In contrast, viral genes expressed later were structural proteins and DNA packaging materials. Transcriptional changes in host genes in response to infection suggested metabolic reprogramming to accommodate the production of virus factories and disruption of host metabolism and development. The transcriptional data from this study expands and improves the TetV-1 genome annotation while highlighting that many of the highly expressed viral genes have unknown functions. These would be good candidates for functional characterization to further improve our understanding of how giant viruses commandeer the host metabolic machinery and facilitate their replication.

## Materials and Methods

### Virus and Algal culture

The virus Tetraselmis virus 1, or TetV-1 for short ^16^, was isolated from Kāneʻohe Bay (21.42973° N, 157.7919° W), a coastal embayment on the northeast shore of Oʻahu, Hawaiʻi that is protected by a barrier reef. A unicellular green alga *Tetraselmis* sp. AL-FL03 (UHM1300) isolated from open ocean waters of Station ALOHA (22.75° N, 158.00° W) in the North Pacific Subtropical Gyre was the initial host used to isolate and propagate the virus, but other strains of *Tetraselmis* isolated from Station ALOHA and Kāneʻohe Bay are susceptible and permissive to the virus ^16^. In this study, a strain from Kāneʻohe Bay, *Tetraselmis* sp. KB-FL45 (UHM1315) was used for the experiments. The uni-algal (but not axenic) culture was maintained in an f/2 medium ^74,75^ at 24 °C on a cycle of 12 h light:12 h dark with a photon flux of 250-280 µE m^-2^ s^-1^ during the light period (07:30–19:30 h). The virus was recently assigned to a new genus and species, *Oceanusvirus kaneohense*, within the family Allomimiviridae ^76^.

### Flow cytometry counts

Algal cells and virions were counted by flow cytometry (Attune^TM^ NxT; Thermo Fisher Scientific) using side scatter (SSC) and either chlorophyll autofluorescence for algae or DNA fluorescence after staining with SYBR Green I (Thermo Fisher Scientific) for the virions to discriminate populations. Autoclaved, filtered seawater was used as a control for the flow cytometer measurements. TetV-1 counting followed the previously published protocol using green fluorescence and side-scatter ^77^.

### Latent period and burst size estimation

For latent period determination, triplicate algal cultures in the exponential growth phase (238 mL each at 1.2 × 10^5^ cells mL^-1^) were each inoculated 4.5 h into the light period with fresh lysate (∼8 mL at ∼1.8 × 10^8^ virions mL^-1^) that had been filtered (0.45 µm Sterivex filter) to remove cell debris, then swirled gently to mix. This resulted in a multiplicity of infection (m.o.i.) of approximately 50. After inoculation, cultures were subsampled periodically up to 30 hours post infection. The cultures, both infected and control, were mixed by swirling gently before sampling. Subsamples (1 mL) were preserved with glutaraldehyde (0.5% v:v final conc.), flash frozen in liquid nitrogen, then stored at –80 °C. Samples were thawed immediately prior to flow cytometric analysis, as described above. Burst size estimates (viruses produced per cell) were calculated by dividing the change in viral counts before (15 hr) and after (30 hr) latent period by the host counts at the beginning of infection. We made the limiting assumption that all cells contributed to viral production because cell debris and clumping at the end of the latent period precluded accurate quantification of any residual unlysed cells. The burst size estimate may, therefore, be somewhat underestimated.

### Experiment for ultrastructural and transcriptional analysis

Guided by data from the first experiment, a second infection experiment was conducted to analyze changes in ultrastructure and transcriptional activity through the latent period. Four independent algal cultures (1350 mL each) were initiated (three experimental and one uninfected control) and grown to exponential phase (approximately 1.48 × 10^5^ cells mL^-1^) under the routine conditions noted above. Approximately 100 mL of filtered virus lysate at ∼1.0 × 10^8^ virions mL^-1^ was added to the three experimental cultures at noon (designated as t = 0 h) for an m.o.i of about 50 as above, and incubations continued for another 20 h. Sub-samples were collected from each culture at t = –1, 0.25, 4, 8, 12, 16, and 20 hours post-infection (hpi) for analyses as follows. One milliliter (1 mL) of the sub-sample was fixed with glutaraldehyde, frozen, and stored as described above for flow cytometry. Fifty milliliters (50 mL) of the subsample were preserved and processed for electron microscopy, as described below. Another 50 mL was immediately filtered through a 2.0 µm polycarbonate filter by vacuum filtration to collect the cells. The filters were transferred to a 2 mL cryovial, flash frozen, and stored at –80°C for subsequent RNA extraction, as described below.

### Transmission electron microscopy (TEM)

Samples for electron microscopy were preserved with a fixative (final concentration: 0.1 M sodium cacodylate, 2% glutaraldehyde, 0.005 M CaCl_2_, 0.06 g glucose/ml). Cells were pelleted at 2,400 × g and embedded in ∼3.3% agarose gel. The embedded samples were washed twice with 0.1 M cacodylate buffer with 0.44 M sucrose for 20–30 minutes. The samples were then post-fixed with 1% osmium tetroxide in 0.1 M cacodylate buffer for 1 hr. Post-fixed samples were dehydrated twice in a graded ethanol series (30%, 50%, 70%, 85%, 95%) for each dilution and thrice in 100% ethanol for 10 minutes each. Samples were then substituted with propylene oxide thrice for 10 minutes each and then infiltrated with 1:1 propylene oxide:epoxy resin (LX112) overnight. They were then immersed in a freshly made 100% epoxy resin for 2 hours, followed by another exchange for 3 hours. Finally, the samples were placed in molds with epoxy resin and polymerized at 60 °C for 2–3 days. Ultrathin (60–80 nm) sections were generated on an RMC MT6000-XL ultramicrotome, viewed unstained on a Hitachi HT7700 TEM at 100 kV, and photographed with an AMT XR-41B 2k x 2k CCD camera. The EM image contrast was adjusted for better clarity.

### RNA extraction and sequencing

Total RNA was extracted from cells collected on filters using a Zymo Direct-zol RNA miniprep kit following the manufacturer’s protocol with the addition of bead beating with ZR bashing beads at the TRIzol lysis step. RNA integrity was checked with gel electrophoresis. RNA concentration was determined by fluorescence using a Qubit RNA High Sensitivity kit (Thermo Fisher Scientific). Quality control using RNA ScreenTape, library preparations, poly-A selection, and Illumina sequencing (HiSeq 2×150 bp) were performed by a commercial facility (Azenta Life Sciences, NJ, USA). The triplicate infected samples at each time point were pooled equally into one sequencing library. This translates to twelve sequencing libraries with one experimental and one control library for each of six time points (–1, 0.25, 4–, 8–, 12– and 16 hours post-infection).

### Virus transcriptome assembly and analysis

Low quality sequencing reads were removed. Adapters and low quality bases were trimmed from the ends of remaining sequences with Trimommatic ^78^ using the following parameters (IlluminaClip:TruSeq3-PE-2.fa:2:30:10:1:true Leading:30 Trailing:30 SlidingWindow:4:15 MinLen:30) and checked with FastQC ^79^. Genome-guided transcriptome assembly was implemented using the Tuxedo protocol (HISAT2, StringTie and Ballgown) ^80–82^. Briefly, the RNA-seq paired-end reads were aligned to the TetV-1 genome using HISAT2. StringTie assembled and quantified expressed genes and transcripts and merged all transcripts from all samples to create a reference set. These Tuxedo-based transcripts were compared to the TetV-1 reference annotation based on Prodigal ^16^. The relative abundance of transcripts calculated as fragments per kilobase of transcripts per million mapped reads (FPKM) were extracted using Ballgown.

### Host transcriptome assembly and analysis

The genome of *Tetraselmis* sp. KB-FL45 has not been sequenced, so a *de novo* transcriptome assembly approach was implemented using Trinity ^83^ after removal of TetV-1 reads using Bowtie2 ^84^. To assess the completeness of the transcriptome, BUSCO was used to detect Chlorophyta single copy genes ^85^. To ensure quality of transcripts, only those that passed TransRate metrics were retained for further analysis ^86^. TransRate evaluates de novo assemblies based on read mapping evidence to detect chimeras, structural and base errors, and incomplete assemblies. Transcript and gene quantification were done using RSEM (RNA-seq by Expectation-Maximization) ^87^, whereas differential expression analysis was performed using DESeq2 in Trinity ^88^. Coding sequence identification and functional annotation were performed using TransDecoder and Trinotate ^89^, respectively, using the following databases: UniProt SwissProt ^90^ (Feb. 2021 release; accessed 04/09/21), Pfam ^91^ (v34.0; accessed 04/09/21), Gene Ontology ^92,93^ (Feb. 2021 release; accessed 04/09/21), Eggnog terms ^94^ (v4.5; accessed 04/09/21). Finally, to assess the enriched functional categories in the differentially expressed gene sets, gene ontology enrichment analysis was performed using topGO ^95^.

## Acknowledgments

This project is funded by the US National Science Foundation to KFE and GFS (OCE #1559356 and OIA #1736030), and a Simons Foundation Early Career Award to KFE. We are indebted to Tina Carvalho of the Biological Electron Microscope Facility, Pacific Bioscience Research Center, for her assistance with the transmission electron microscopy. The computational work in this paper was implemented at the University of Hawai‘i High Performance Computer. The technical support and advanced computing resources from the University of Hawaii Information Technology Services – Cyberinfrastructure, funded partly by the National Science Foundation MRI award # 1920304, are gratefully acknowledged. We thank Feresa Cabrera for assisting in one of the time series. We are also grateful to Rosie Alegado for valuable suggestions on improving this manuscript. This work is supported by the Hawai‘i Institute of Geophysics and Planetology Denise B. Evans Fellowship, CMAIKI (Center for Microbiome Analysis and Investigation Graduate Research Competition Fund, UH Mānoa) and Uehiro Center for the Advancement of Oceanography Graduate Student Fellowship to APG.

## Data Accessibility

The raw sequencing reads used in this paper have been deposited in the NCBI Short Read Archive as BioProject PRJNA1102674. Whereas the codes used in this study are deposited in GitHub (https://github.com/andriangajigan/TetV_RNA/). The host transcriptome assembly, annotation, and differential expression data are available at FigShare (https://figshare.com/s/98d2742f5f1c1f3bf6fb, and https://figshare.com/s/aad55ae781891e083b09). All other data are available as supplementary material.

## Author Contributions

APG, CRS, and GFS conceived of the paper. APG and CRS did the experiments. All authors (APG, CRS, CC, KFE, GFS) contributed to the data analysis, writing, and approving the manuscript.

