## Supplementary material for "Ultrastructural and transcriptional changes during a giant virus infection of a green alga": Supp Figure 1-9, Supp Table 2-3

### Supplementary Figures and Tables

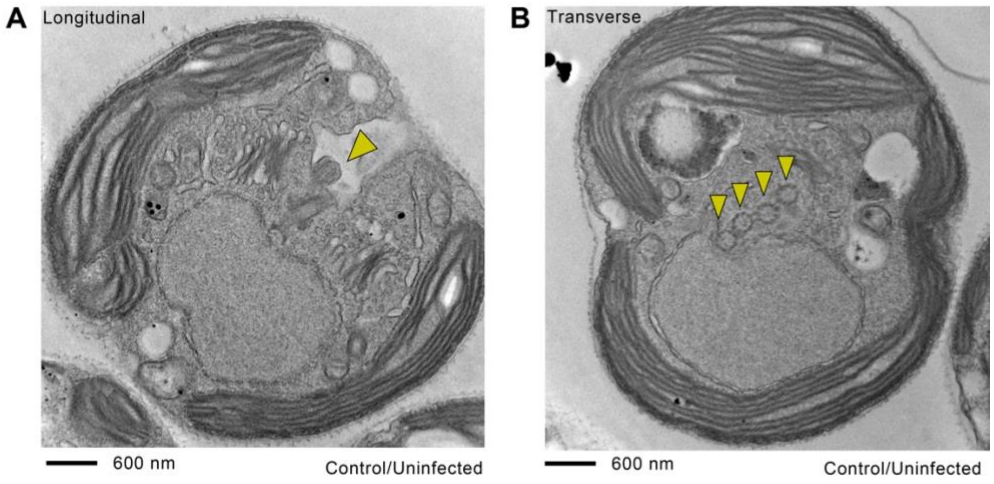


**Supplementary Figure 1. *Tetraselmis* sp. KB-FL45 basal bodies.** Electron micrographs showing **(A)** longitudinal and **(B)** transverse cross sections of *Tetraselmis* sp. Arrows indicate the four basal bodies (base of flagella).

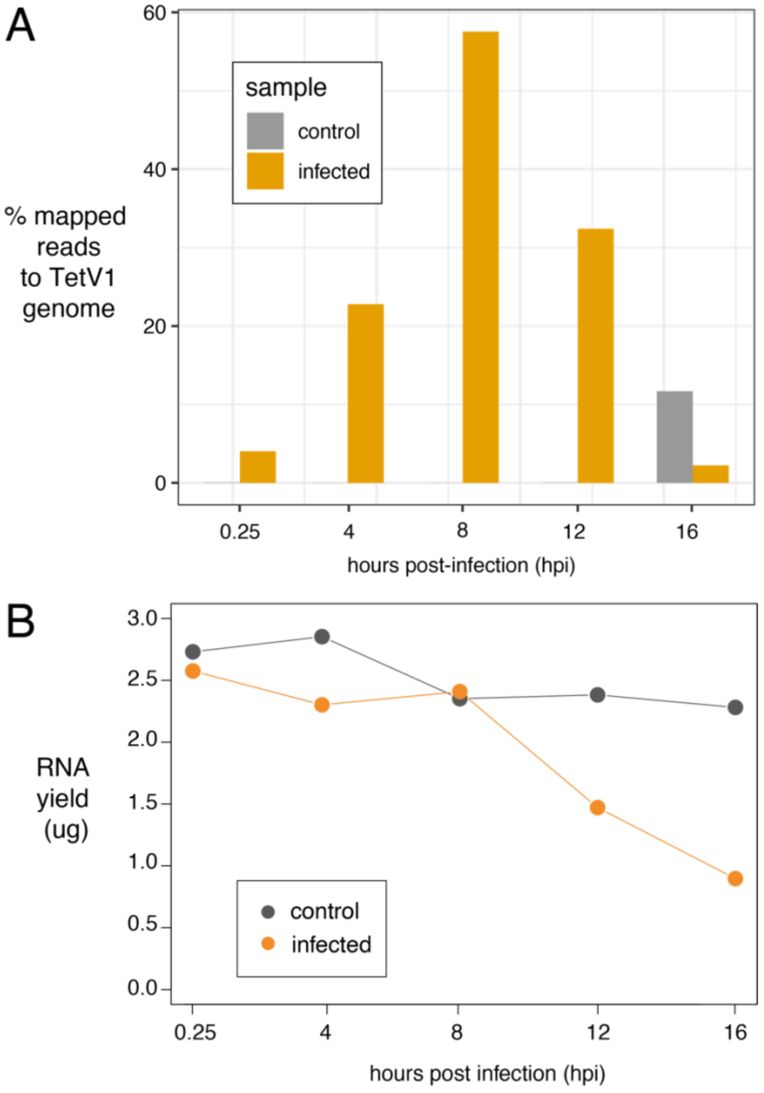


**Supplementary Figure 2. RNA-seq mapping and RNA yield. (A)** High viral RNA reads coincide with the increasing presence of intracellular virions. As virus reads were detected in the control sample at 16 hours, this sample was used only for host genome assembly after removal of virus reads and was not included in gene expression analysis. Thus, 0.25 and 4hpi were used as early-stage samples while 8 and 12hpi represents the late-stage samples for host gene expression analysis **(B)** RNA yield remained the same for controls but decreased in infected cultures over time.

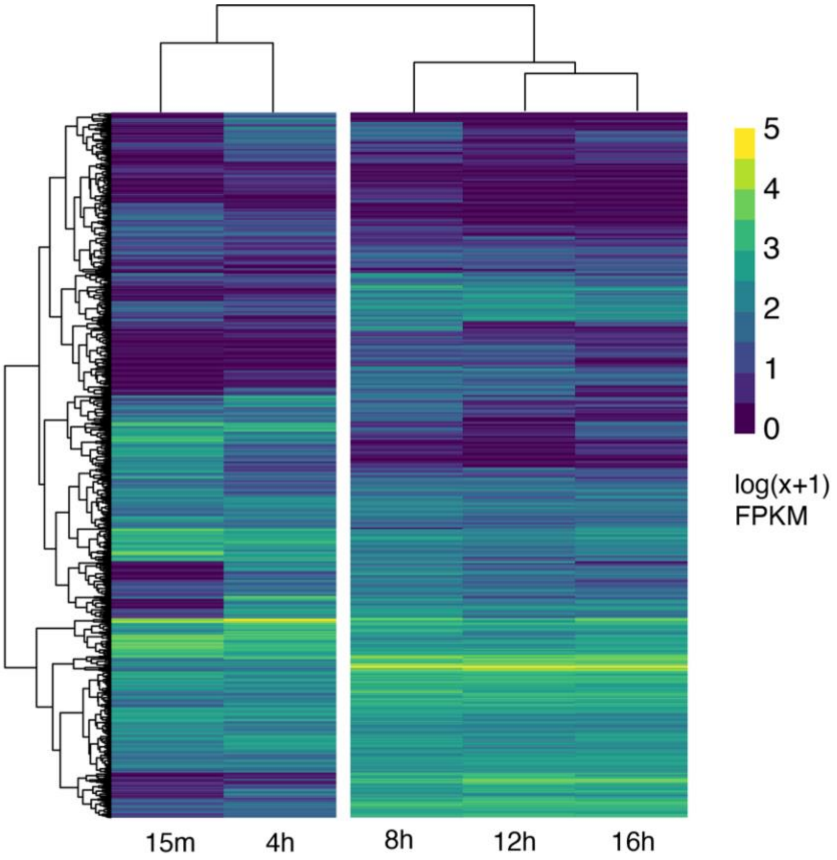


**Supplementary Figure 3. Global TetV-1 gene expression.** Time points cluster together based on global gene expression patterns. FPKM = Fragments Per Kilobase of transcript per Million mapped reads.

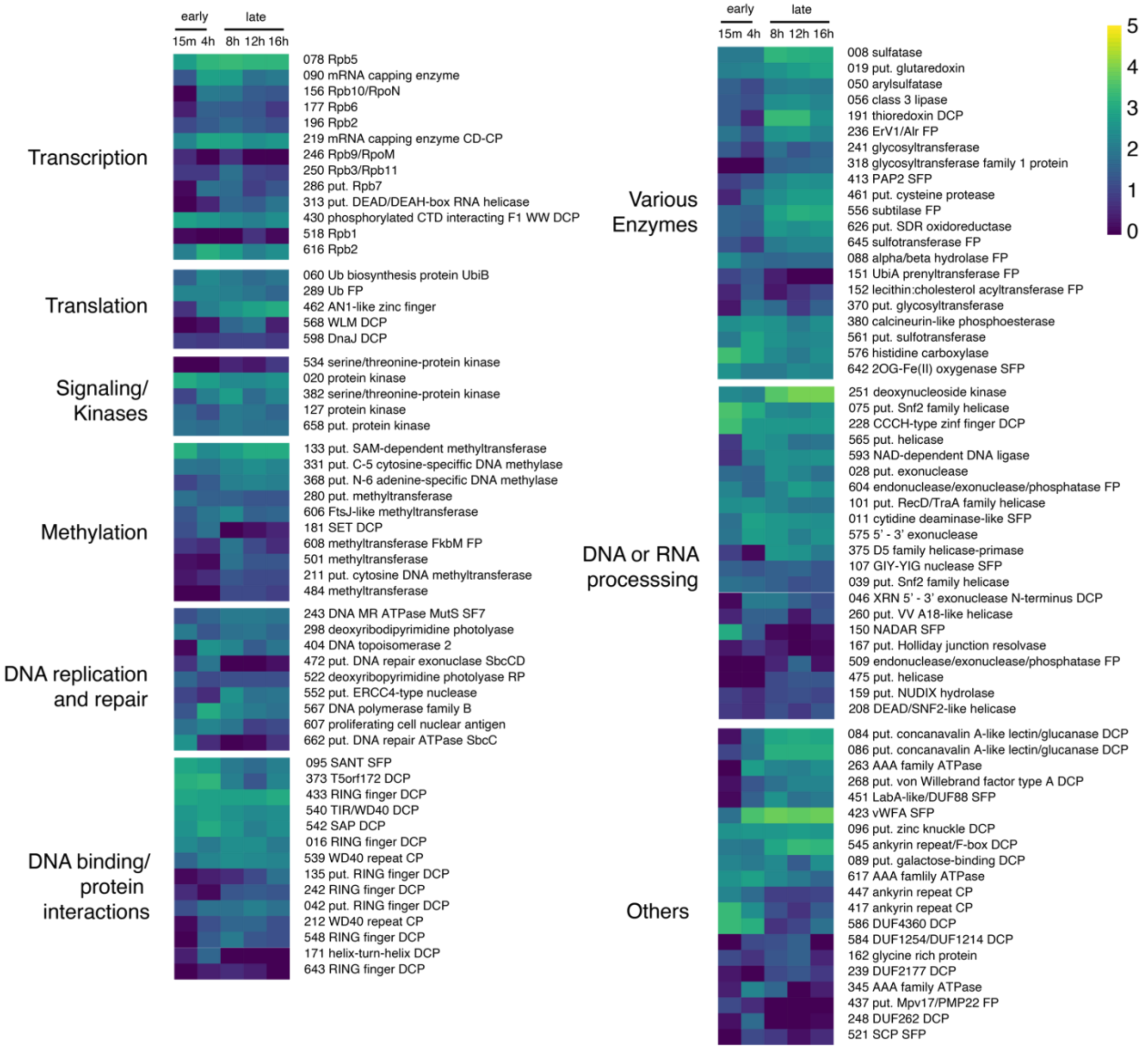


**Supplementary Figure 4. Temporal expression of select TetV1 genes.** Put. = putative, Ub = ubiquitin, DCP = domain-containing protein, SFP = superfamily containing protein, FP =family protein, CD-CP = catalytic domain-containing protein, FPKM = Fragments Per Kilobase of transcript per Million mapped reads.

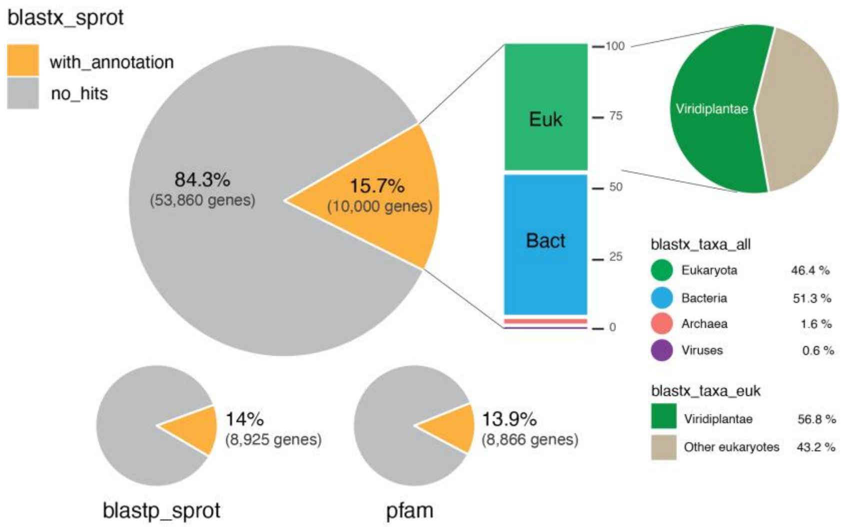


**Supplementary Figure 5. *Tetraselmis* sp. KB FL45 functional annotations.** BLAST matches to SwissProt and Pfam annotations.

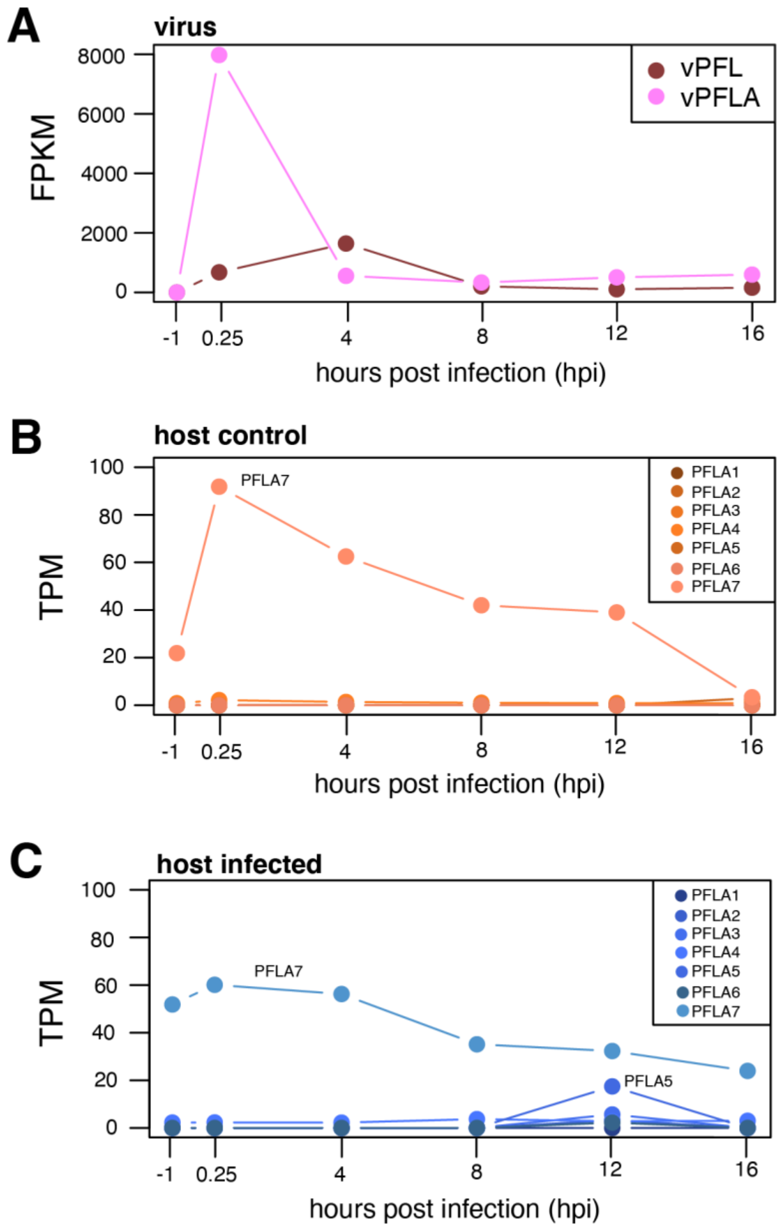


**Supplementary Figure 6. Temporal expression of fermentation-associated genes. (A)** Virus encoded pyruvate formate lyase (vPFL) and pyruvate formate-lyase activating enzyme (vPFLA). Temporal expression of seven host pyruvate formate-lyase (PFL) in **(B)** control time course and **(C)** infected time course. Host PFL is not detected in the host transcriptome. FPKM = fragment per kilobase transcript per million mapped reads; TPM = transcript per million.

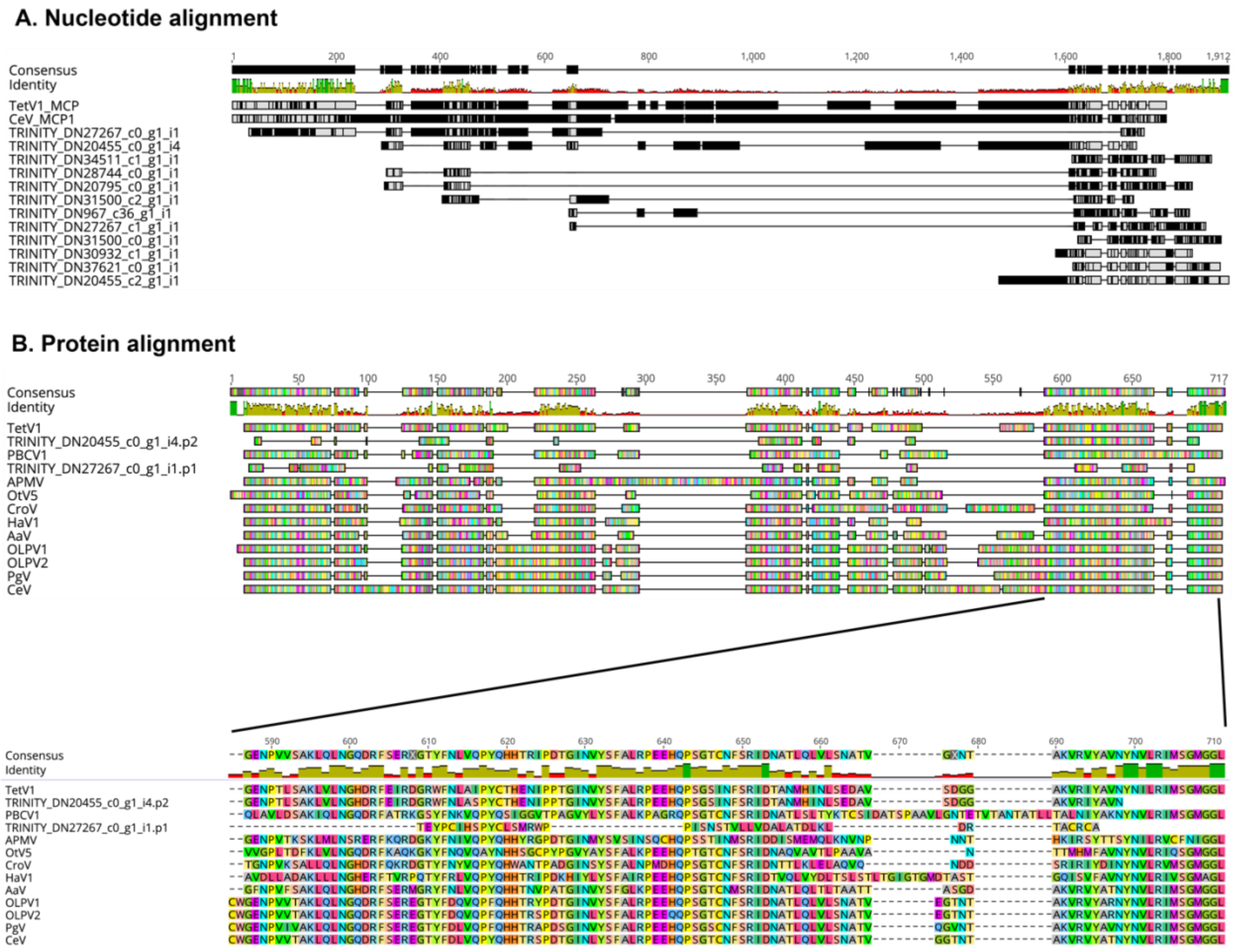


**Supplementary Figure 7. Putative endogenized major capsid proteins.** (A) Nucleotide alignment of putative endogenized MCPs. (B) Protein alignment of two putative endogenized MCPs (TRINITY_DN20455_c0_g1 & TRINITY_DN27267_c0_g1) that were upregulated and passed the TransDecoder cutoff of >100 amino acids. Alignment implemented using MUSCLE 3.8 [96] in Geneious v11.1. Figures generated using Geneious v11.1

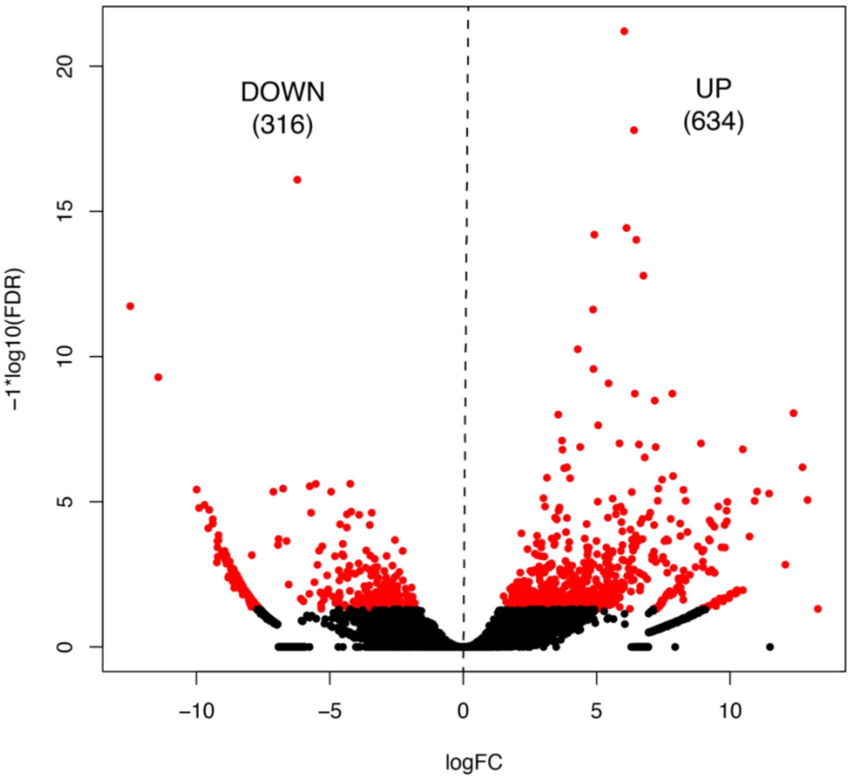


**Supplementary Figure 8. Differential expressed genes in *Tetraselmis* sp. between day and night cycle.** Volcano plot showing upregulated and downregulated genes in day (n=3) vs night (n=2) comparisons at FDR/adjusted p-value ≤ 0.05 (red). Numbers in the parentheses are number of genes in each group. FC = fold change; FDR = false discovery rate/adjusted p-value.

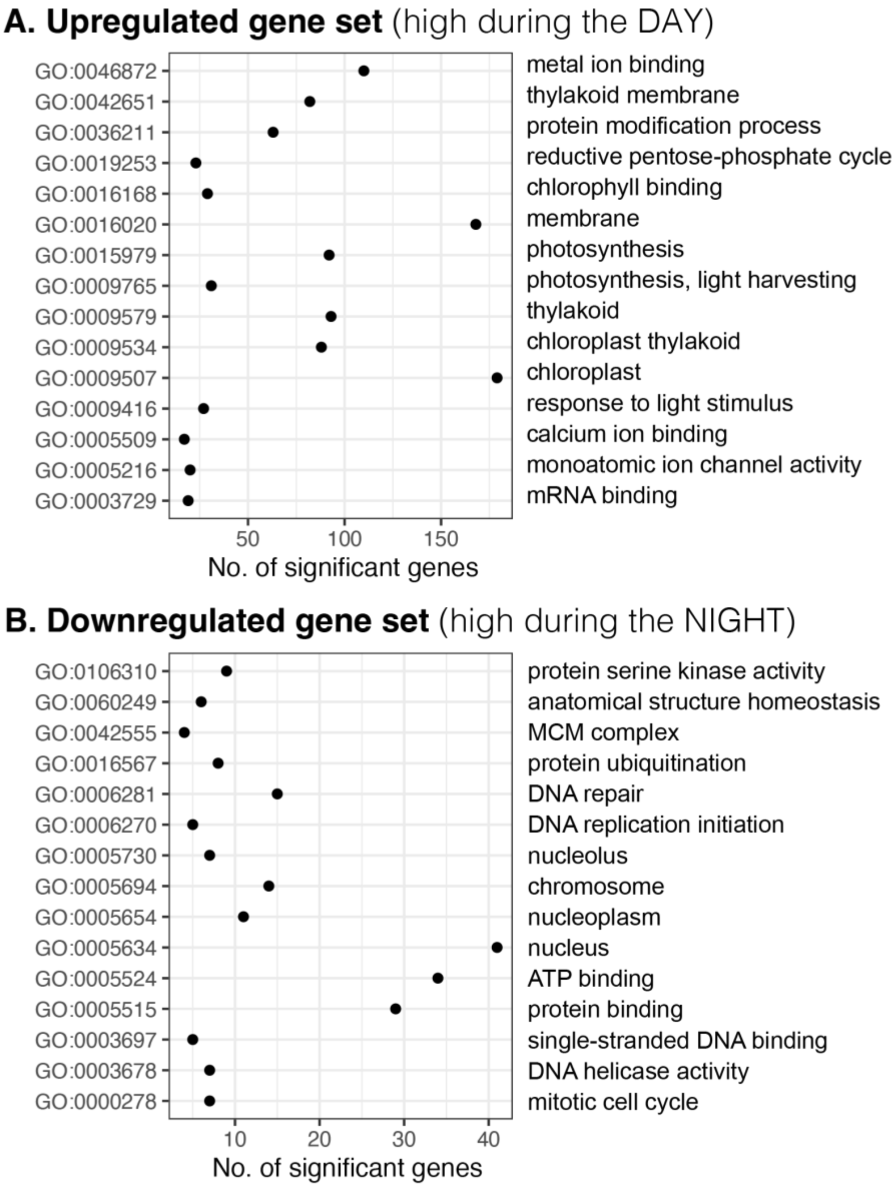


**Supplementary Figure 9. Gene enrichment analysis for light vs dark cycle.**
All enriched GOs shown in the graphs have p-values <<< 0.05

**Supplementary Table 2. Trinity *de novo* assembly metrics and statistics for *Tetraselmis* sp. KB FL45**

| Parameter | **Tet_all** (using all samples, no virus reads) | **Tet_host** (using only the uninfected control) | ***T. striata*** (gene model) [a] |
|---|---|---|---|
| Libraries used | 12 | 6 | - |
| Genes | 69,817 | 44,046 | 46,696 |
| Transcripts | 159,006 | 116,016 | - |
| GC% | 53.77 | 54.36 | 65.05 |
| N50 | 3,244 | 3,015 | 1,725 |
| % Raw read alignment | 98.84 | 98.97 | - |
| BUSCO [b] | 96% | 95% | - |

[a] Using the Tetstr1_GeneCatalog_transcripts_20200803.nt.fasta (manually curated best gene models in PhycoCosm)
[b] Percent of Chlorophyta single copy genes found in the assembly (out of 1519 genes)

**Supplementary Table 3. Filtering steps and metrics for *Tetraselmis* sp. KB FL45 transcriptome**

| | **Filtering step** | **genes** | **transcripts** | **%GC** | **N50** | **BUSCO* completeness** |
|---|---|---|---|---|---|---|
| 1 | Trinity raw output (Tet_all, no virus reads) | 69,817 | 159,006 | 53.8 | 3,244 | 95.7% (74.7% duplicated) |
| 2 | TransRate (retain only good.fasta) | 63,860 | 137,936 | 53.9 | 3,081 | 95.5% (72.9% duplicated) |
| 3 | Highest expressed isoform per gene | 63,860 | 63,860 | 53.4 | 1,801 | - |

*using Chlorophyta single copy genes (Chlorophyta_odb10)
